# Response of microbial community function to fluctuating geochemical conditions within a legacy radioactive waste trench environment

**DOI:** 10.1101/121376

**Authors:** Xabier Vázquez-Campos, Andrew S. Kinsela, Mark W. Bligh, Jennifer J. Harrison, Timothy E. Payne, T. David Waite

## Abstract

During the 1960s, small quantities of radioactive materials were co-disposed with chemical waste at the Little Forest Legacy Site (Sydney, Australia) in three-metre-deep, unlined trenches. Chemical and microbial analyses, including functional and taxonomic information derived from shotgun metagenomics, were collected across a six-week period immediately after a prolonged rainfall event to assess how changing water levels impact upon the microbial ecology and contaminant mobility. Collectively, results demonstrated that oxygen-laden rainwater rapidly altered the redox balance in the trench water, strongly impacting microbial functioning as well as the radiochemistry. Two contaminants of concern, plutonium and americium, were shown to transition from solid-iron-associated species immediately after the initial rainwater pulse, to progressively more soluble moieties as reducing conditions were enhanced. Functional metagenomics revealed the potentially important role that the taxonomically-diverse microbial community played in this transition. In particular, aerobes dominated in the first day followed by an increase of facultative anaerobes/denitrifiers at day four. Towards the mid-end of the sampling period, the functional and taxonomic profiles depicted an anaerobic community distinguished by a higher representation of dissimilatory sulfate reduction and methanogenesis pathways. Our results have important implications to similar near-surface environmental systems in which redox cycling occurs.

**Importance:** The role of chemical and microbiological factors in mediating the biogeochemistry of groundwaters from trenches used to dispose of radioactive materials during the 1960s is examined in this study. Specifically, chemical and microbial analyses, including functional and taxonomic information derived from shotgun metagenomics, were collected across a six-week period immediately after a prolonged rainfall event to assess how changing water levels influence microbial ecology and contaminant mobility.

Results demonstrate that oxygen-laden rainwater rapidly altered the redox balance in the trench water, strongly impacting microbial functioning as well as the radiochemistry. Two contaminants of concern, plutonium and americium, were shown to transition from solidiron-associated species immediately after the initial rainwater pulse, to progressively more soluble moieties as reducing conditions were enhanced. Functional metagenomics revealed the important role that the taxonomically-diverse microbial community played in this transition. Our results have important implications to similar near-surface environmental systems in which redox cycling occurs.

## Introduction

The rapid expansion of an emerging nuclear industry immediately following World War II resulted in substantial volumes of low-level radioactive waste (LLRW) being generated from nuclear fuel cycle, weapons production, medical radioisotope and radiochemical research activities. Although there was no consensus at this time, low-level waste (and in some cases more active material) was commonly disposed of by burial in shallow trenches, as evidenced in the United States at Maxey Flats (1), Oak Ridge (2) and Hanford (3, 4); Canada at Chalk River (5); in the United Kingdom at Harwell (6) and LLRW (7); in Lithuania at Maišiagala (8); and more recently in Ukraine at Chernobyl (9) to name but a few. This was also the case for Australia’s only nuclear (research) reactor at Lucas Heights. Known as the Little Forest Legacy Site (LFLS), radioactive materials, including minor amounts of ^239+240^Pu and ^241^Am were placed in narrow (0.6 m), 3 m deep, unlined trenches from 1960 to 1968 (10–12). Large volumes of contaminated non-radioactive materials and equipment were also disposed of in these trenches (10).

The LFLS trenches were excavated within undisturbed geological matrices of red-brown and grey clay (primarily kaolinite and illite/smectite) derived from the underlying weathered shale, interspersed with minor phases of hematite and goethite (11, 13). The site was chosen in part due to the low hydraulic conductivities of these materials (~ 9 to 66 mm/day) in order to isolate the LLRW from waters associated with the local hydrology cycle (10–12). However, an unintended consequence of this (also experienced at other disposal sites) has been that periodic intense rainfall and prolonged dry conditions can facilitate complete saturation and desaturation of the more permeable waste material via surface infiltration and evapo(transpi)ration/leakage mechanisms, respectively. Elaborating further, this frequently results in infiltrating water filling up the more porous trenches to the surface (akin to a ‘bathtub’), which has been shown to be a primary mechanism for the dispersion of contaminants ^239+240^Pu and ^241^Am (10). Furthermore, this has allowed for redox cycling to occur unabated in the LLRW trenches since their construction, potentially promoting redox tolerant plasticity in the microbial communities present (14).

With actinide mobility, in many instances, strongly dependent on their oxidation state (15), microbial communities can potentially play decisive roles in determining the fate and mobility of such elements in the environment. In general, microbial communities can influence actinide chemistry by partaking in redox (16–18), dissolution (19, 20), precipitation (21, 22), sorption (23) and/or methylation (24) reactions, which may either enhance or retard contaminant mobility.

Despite this, much of the scientific research to date has focussed on isolated individual species (such as *Geobacter*, *Shewanella* and *Clostridium*) and their impact on actinide, particularly uranium, behaviour. The challenge remains as to how best to identify the role of microorganisms and their functioning within the wider microbial community which may be undergoing external, environmental changes (25). Furthermore, subsurface environments such as aquifers and shallow groundwaters, aside from being poorly studied, have been shown to be havens for microbial novelty not dominated by the well-characterised organisms listed above (26). Culture-independent techniques, such as metagenomics, in which genomic sequences capture the aggregate microbial ecology of a sample (27, 28) have the potential to enhance our understanding of these complex biogeochemical systems.

The question remains at LFLS, and in similar contaminated redox-cycling environments, as to what function microbial communities may be performing in the direct or indirect mobilisation and/or retention of legacy radionuclides. As the establishment of causality between the legacy contaminants and microbial communities cannot be achieved in such a non-invasive environmental study, the aim of this research was to understand the role that periodic inundation from a large rainfall event, and presumably oxygen penetration, had on the concomitant changes to chemistry-radiochemistry and microbial communities in a LLRW trench environment. The two contaminants of major concern, Pu and Am, were the focus of this research with shotgun metagenomics used to examine the microbial systems function and taxonomy.

## Materials and methods

### Sample collection

Trench water samples were collected on five separate occasions across a two-month period from LFLS. They were obtained from a screened PVC pipe which extends 1.55 m below the ground surface. This pipe provides the only point of access into the legacy trenches and was opportunistically installed during the partial collapse of the trench surface. Further details of the trench sampler have been previously described in the literature (10, 29). Briefly, a peristaltic pump operating under low flow rate was used to purge the borehole until chemical parameters, particularly pH and oxidation/reduction potential (ORP), became stable (Supplementary File 1). Measurements were made using a multi-probe system (YSI 556 MPS) coupled to a 100 mL flow cell. Stability usually took between 30 and 60 min, after which sample collection began. Chemical parameters, along with the water depth, were monitored over the sampling period (typically some hours) on the day of sampling to ensure the water was representative of that particular period. The stability of flow cell parameters and the fact that the water level did not decrease during the collection of 4–5 L of sample material provides assurance that the samples were representative of the particular event rather than anomalous local conditions.

The sample collection period commenced (on 23^rd^ April 2015) immediately after a substantial rainfall event, during which ~220 mm of rain fell over 3 days (Figure S1). Trench water samples were subsequently collected on the 27^th^ and 29^th^ April, 14^th^ May and 9^th^ June (or days 0, 4, 6, 21 and 47, post-rain, Figure S1).

Samples for chemical analyses were collected both with and without an in-line filtration membrane (0.45 μm PES, Waterra FHT-Groundwater) in pre-washed and trench water rinsed HDPE bottles except samples for organic carbon analysis which were collected in glass bottles. Samples for radiochemical and cation analyses were acidified using double-distilled HNO_3_. All samples were then stored at 4 °C until analysis.

Samples for microbial analyses were collected by directly filtering the trench water using a sterile PES 0.22 μm syringe filter (Sterivex-GP, Merck Millipore) with the volumes of filtrate passing through the membranes recorded (410.1±82.2 mL per replicate, mean ± SD). Filters were capped on site before being stored at -18 °C until analysis. Microbial sampling was performed in triplicate to assess inherent variation.

Due to regulatory protocols governing sampling at the site, no solid phase material could be extracted from the trenches. As the surrounding geological materials into which the trenches were excavated provide little resemblance both physically and chemically to the deposited waste within the trenches, they were discarded as providing useful information for this study. As such all sampling within the trenches pertains to the extraction of soluble or suspended solid-phases from the screened borehole.

### Chemical analyses of trench waters

The primary radiochemical contaminants, ^239+240^Pu and ^241^Am, were separated from other radiochemical fractions using TEVA^™^ and TRU^™^ resin cartridges (30) before measuring activity by alpha spectroscopy using a Canberra Alpha Analyst system coupled with Passive Implanted Planar Silicon detectors (Canberra).

Cations (including Si, S and P) were measured by either ICP-AES or ICP-MS depending on relative concentration. Anions (F^-^, Cl^-^, Br^-^, I^-^, PO_4_^3-^ and NO_3_^-^) were determined by ion chromatography (Dionex DX-600 IC System).

Non-purgeable dissolved organic carbon (0.45 μm filtered) was measured on freshly-collected replicate samples by combustion catalytic oxidation (TOC-5000A, Shimadzu) with concentration determined by a five-point calibration using newly-prepared potassium hydrogen phthalate solutions.

Ferrous iron concentrations from 0.45 μm filtered samples were preserved in the field with ammonium acetate-buffered phenanthroline solution (31). The absorbance of Fe(II) was measured at 510 nm (USB4000, Ocean Optics) calibrated against a freshly-prepared ammonium ferrous sulfate solution.

In field measurements of dissolved sulfide were attempted using methylene blue colorimetry, but were consistently below detection limits (~1 μM) at all sampling time points.

All chemical analyses were performed in triplicate.

### DNA extraction and sequencing

The PowerLyzer^®^ PowerSoil^®^ DNA Isolation kit (MO BIO) was used to extract DNA from Sterivex filters following the modified protocol of Jacobs et al (32). Bead beating was performed singularly in a PowerLyzer^®^ 24 Bench Top Bead-Based Homogenizer (MO BIO) at 2000 rpm for 5 min.

Concentration and DNA quality was evaluated with LabChip^®^ GX (PerkinElmer). Libraries were prepared with the TruSeq Nano DNA Library Preparation Kit and sequenced in a NextSeq 500 (Illumina) with a 2×150 bp high output run. Output consisted of paired-end reads with a median insert size of ~480 nt. Reads were deposited at the European Nucleotide Archive (PRJEB14718/ERG009353).

### Data processing and functional analyses

Raw sequencing reads were pre-processed with Trim Galore! (http://www.bioinformatics.babraham.ac.uk/projects/trim_galore/) to remove residual adapters as well as short and low quality sequences (Phred33 ≥ 20) while keeping unpaired reads.

Taxonomic profiles were obtained from shotgun metagenomic data by extracting 16S rDNA gene sequences with GraftM v0.10.1 (https://github.com/geronimp/graftM) and mapping them with the 97% clustered GreenGenes database provided by GraftM developers (package 4.39).

Functional analyses of the metagenomic data were performed with HUMAnN2 v0.6.0 (33). Gene families were grouped by MetaCyc reactions (RXN), keeping unmapped and ungrouped reads for calculating copies per million and relative abundances. MetaCyc RXN groups were filtered and only those with at least two replicates from the same sampling time-point with a relative abundance of 10^-4^ or higher were analysed (33). MetaCyc RXN notations have been used as is within Results and Discussion to provide an exact match with the database. RXNs can be considered equivalent to the more traditionally used KO terms from the KEGG database. Grouping by GO terms, specifically performed to evaluate virus presence, indicated values under the threshold levels for all the virus-specific GO terms identified. As such, viruses were not taken in further consideration.

Analysis of variance was performed with the software package STAMP v2.1.3 (34) using the Benjamini-Hochberg False Discovery Rate approach (35) for correction of *p*-values.

## Results and Discussion

### Water level changes and chemical analyses

The initial 220 mm rainfall event resulted in trenches filling to capacity and discharging from the surface or porous near-surface (0–0.2 m) in the ‘bathtub’ mechanism described by Payne et al (10) (Figure S1). Despite the subsequent 47-day sampling period receiving a further 72 mm of rainfall, which periodically increased trench water levels, an overall decline in trench water levels was observed across the sampling period (Figure S1).

Across the sampling period and as the water level declined, the pH was observed to increase from 6.30 to 6.60, whereas the *E_h_* values decreased from 247 to 147 mV (Figure 1). Of the cations and anions analysed (Figure 2), iron displayed one of the greatest variations in concentration, increasing from 0.42 mM at day 0, to 1.00 mM by day 47. The elevated concentrations of iron are unsurprising, given that the LLRW at LFLS was disposed of in-part within ~760 steel drums (36), and buried within a highly weathered shale geological matrix (11). The continuous presence of Fe(II), even at day 0, confers reducing conditions in excess of *E_h_* values recorded, casting uncertainty over the absolute values supplied by the *E_h_* probe. The increase in Fe(II) concentrations with time was likely due to Fe(III) oxyhydroxide reduction, an observation supported by the concurrent liberation of Si and P, two elements that are typically co-associated with Fe(III)-oxyhydroxide at LFLS (37). The constant concentrations of expected conservative elements Cl^-^ and K suggest that the drop in the trench water level was due to subsurface outflow rather than evaporative processes across the course of sampling. Other important elemental transitions included a 10-fold decrease in dissolved sulfur concentrations (presumed to be sulfate), from an initial concentration of 62.4 to 6.2 μM at day 47 (Figure 2). Nitrate concentrations doubled between days 0 and 6 (from 0.24 to 0.55 μM) after which they decreased to below detection limits at days 21 and 47 (Figure 2).

**Figure 1.**
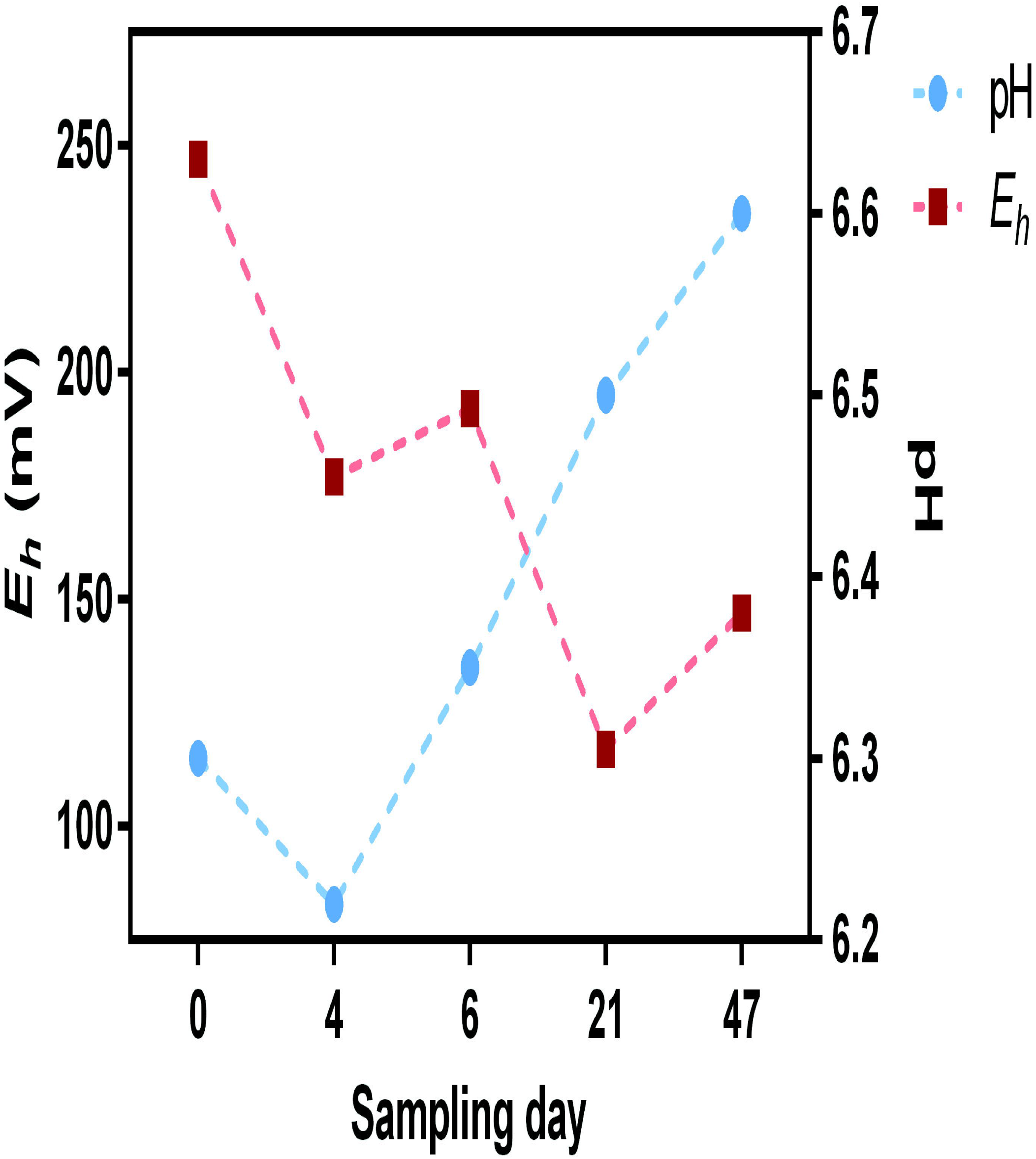
pH and *E_h_* (Standard Hydrogen Electrode corrected) measurements from the trench water across the sampling period.

**Figure 2.**
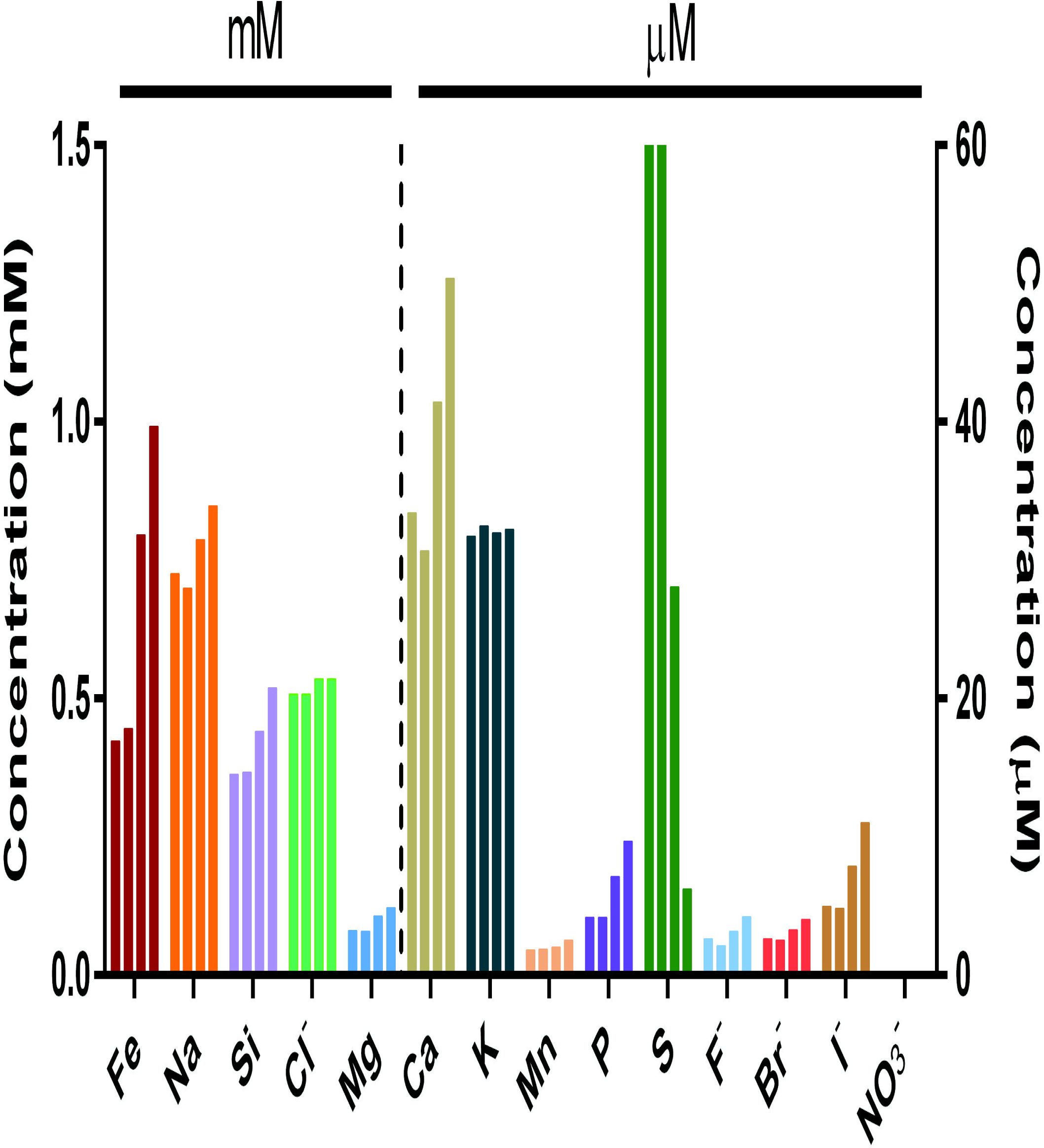
Temporal changes to element/ion concentrations in the trench water. Bars show the individual concentrations of each element/ion over the five sampling days 0, 4, 6, 21 and 47.

The total (unfiltered) ^241^Am activity increased from 15.7 to 27.8 Bq/L (1.8-fold increase) while the soluble (filtered) fraction increased 3.4-times (7.2 to 24.7 Bq/L) between days 0 and 47 (Figure 3). Although the activity of the total ^239+240^Pu showed a proportionally smaller increase than ^241^Am across the sampling period, from 30.4 to 45.6 Bq/L (1.5 times), the soluble fraction increased from 0.21 at day 0 and reached a maximum of 0.80 (35.1 Bq/L) by day 21. Despite the low-flow sampling conditions, the majority of ^241^Am and ^239+240^Pu in the trench water was solid-associated at Day 0. This implies that the solid-associated actinides are relatively stable or easily (re)mobilised, and also that during the rapid influx of rainwater when the trenches are filling up (and potentially overspilling), colloidal transport is likely to be the major form of contaminant movement at this site. Previous redox state measurements at LFLS found that Pu was present in the Pu(IV) (64.5%) and colloidal/Pu(III) (35.5%) states whilst Am was present exclusively as Am(III) (29). The vast quantities of Fe(II) as well as the results of ancillary measurements (DO, ORP) would suggest that Pu(IV) and Am(III) were again the dominant oxidation states of the trench water contaminants. The results of all water quality parameters measured are provided in Table 1.

**Figure 3.**
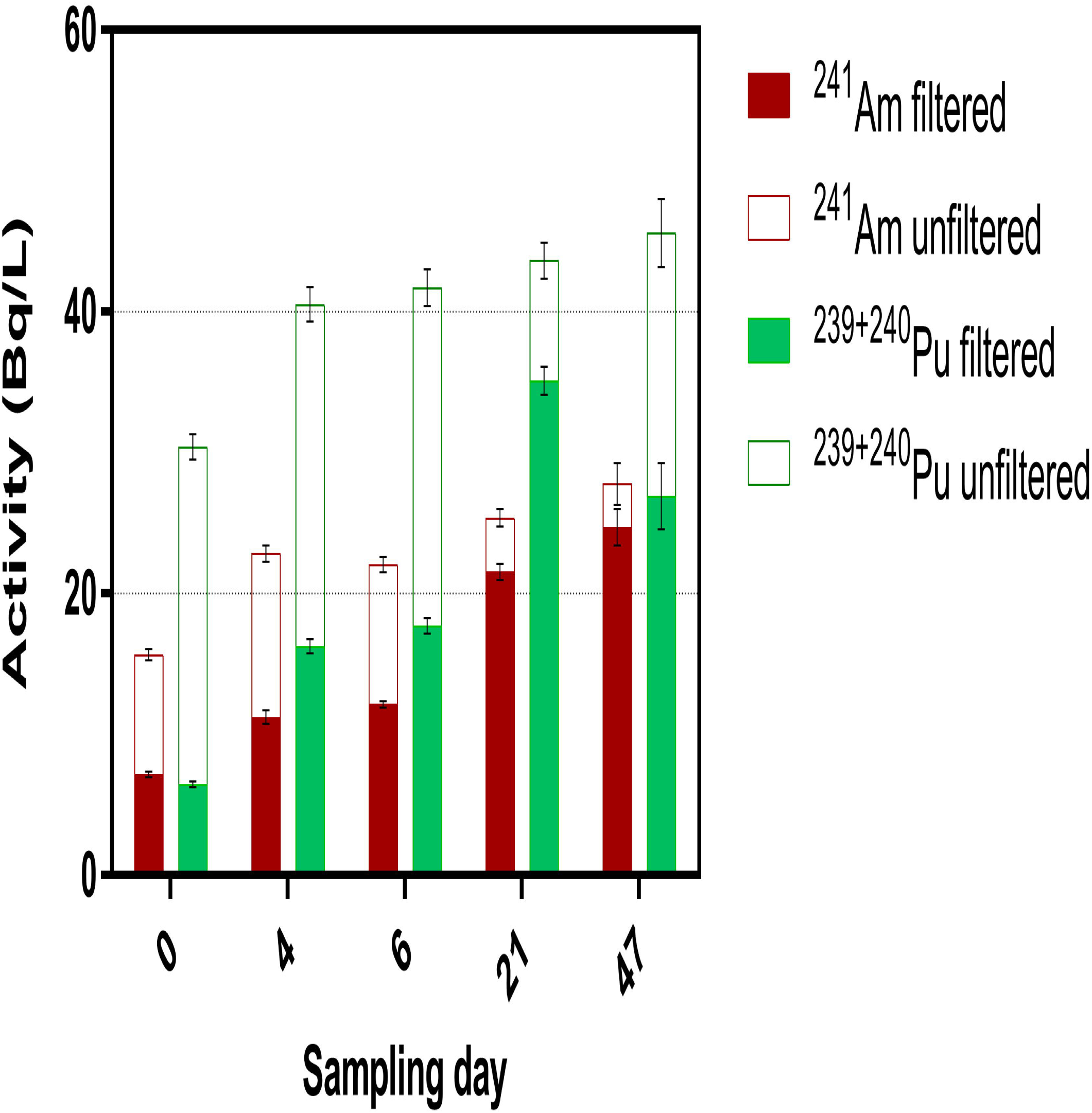
Activity of filtered and unfiltered radionuclides measured in the trench water. Filtered fractions (<0.45 μm, solid fill) are equivalent to soluble and smaller colloidal particles. Unfiltered fractions (full bar) are as considered total (soluble plus all suspended solids) concentrations. Error bars show the standard deviation of triplicate measurements.

### Community composition

Community profiles showed that *Bacteria* dominated over *Archaea* in the trench water across the entire sampling period. A total of 70 phyla out of the 85 included in the GreenGenes database were detected at some point and 11 phyla with values over 1% of the total community at all times. Eukarya were found in low (~1% to ~2%), nearly constant abundance trough the samples. Similarly, eukaryotic-specific RXNs were all below our threshold significance level. As such they are not considered in further detail nor for the taxa abundance numbers throughout the manuscript.

Archaea oscillated between 2.6 and 10.6% of the classified reads with minimum and maximum at days 4 and 47, respectively. *Micrarchaeota* and *Parvarchaeota* (DPANN) were the most abundant phyla at all times with a combined 45.5–55.9% of *Archaea*, while remaining sequences were shared in variable proportions between the TACK superphylum and *Euryarchaeota* (Figure 4). Although the fraction of TACK decreased over time from 24.8% at day 0 to 12.4% at day 47, *Euryarchaeota* reached a maximum at day 47 contributing 37.2% of all *Archaea*. These changes were derived mainly from variations in the SAGMA-X family (*Thaumarchaeota*), related to the ammonia oxidising archaeon ‘*Ca.* Nitrosotalea’ (38, 39) and, in *Bathyarchaeota* (MCG), which includes the only potential non-euryarchaeal methanogens (39, 40) and/or one of the few acetogenic *Archaea* (41). The increase in total *Archaea* abundance over time was related to changes to and ANME-2d, *Methanoregulaceae* and *Methanosaetaceae* families, implicated in methane metabolism (42–44) as well as the *Micrarchaeota* and *Parvarchaeota* phyla.

**Figure 4.**
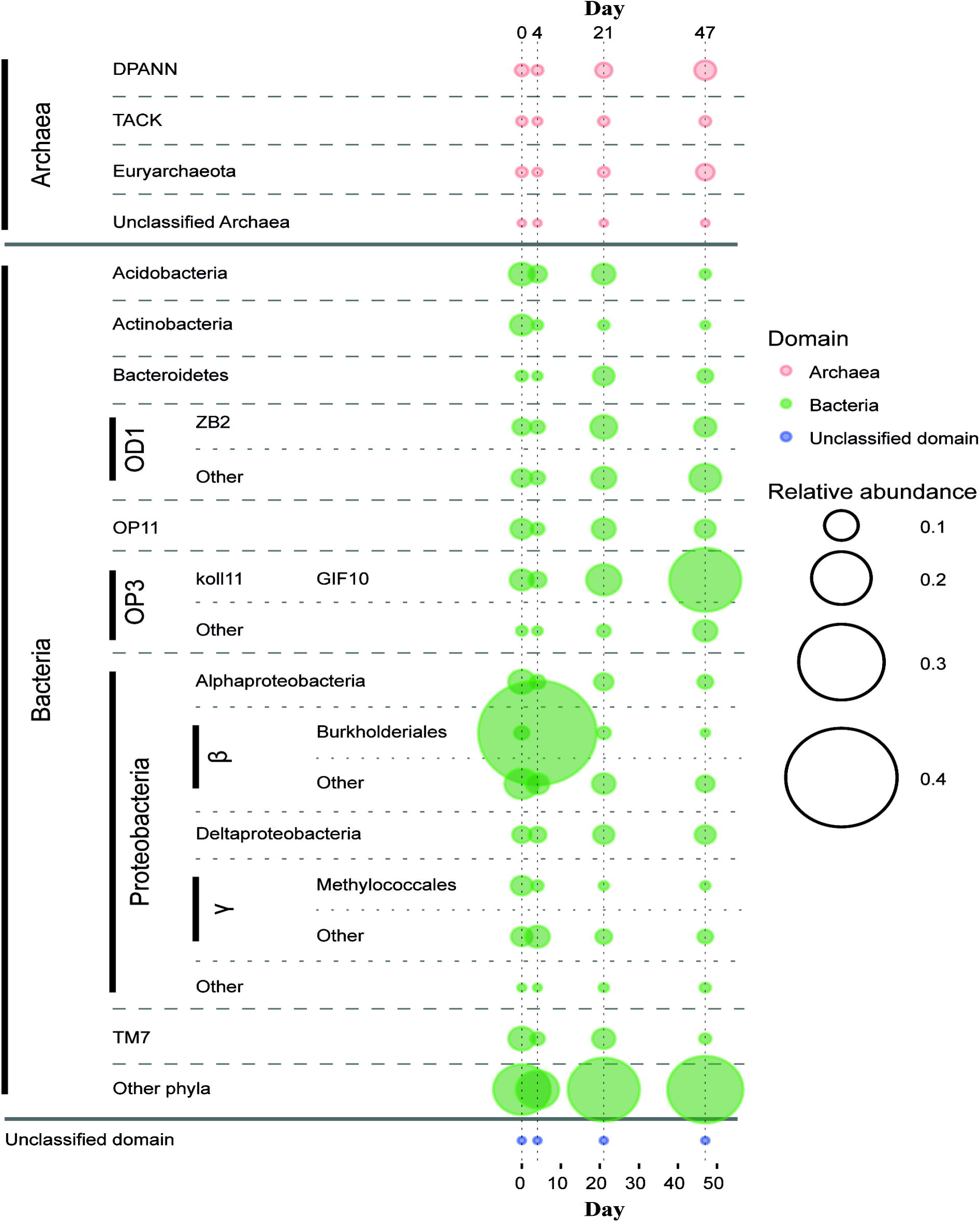
Taxa relative abundances over time. Only taxa with an average relative abundance >5% of the total sequences in at least a single sampling day are shown. Taxa with <5% of all sequences are grouped under ‘other’. Phyla with <5% of relative abundance at all times are grouped under ‘other phyla’, except Archaea.

Only 8 bacterial phyla grouped more than 5% of the total bacterial reads at any one sampling point, and amongst them, only *Chloroflexi*, OD1 (*Parcubacteria*), OP11 (*Microgenomates*), OP3 (*Omnitrophica*) and *Proteobacteria* were within the 10 most abundant bacterial phyla at all times (Figure 4). The principal changes observed over time refer to *Proteobacteria*, TM7 (*Saccharibacteria*), OP3, *Acidobacteria* and OD1. While the proportion of TM7 and *Acidobacteria* decreased from day 0, both OP3, and OD1 reached their maximum by day 47 (18.9% and 14.7%). The *Proteobacteria*, often the most abundant phylum, showed a conspicuous increase, mainly due to *Betaproteobacteria*, comprising 48.7% of all classified reads at day 4. The order *Burkholderiales* was identified as being primarily responsible for this change (42.6% of total), esp. the genus *Ralstonia* (20.3%) within *Oxalobacteraceae* (34.0%). These bacteria were also responsible for the altered functional profile observed at day 4.

Several studies focusing on the microbiology of groundwater or subsurface ecosystems have demonstrated the existence of organisms able to pass through standard 0.22 μm filters (45–47). Despite this, a substantial portion of the Little Forest trench water community corresponded to taxa known to be able to pass through 0.22 μm filters, i.e. OD1, OP11 and DPANN. We suggest this could be derived from the fast precipitation of iron in our extracted trench waters, effectively entrapping cells, or that they were attached to colloidal particles or to other cells. Still, it is possible that the numbers for these ‘ultra-small’ taxa could be underestimated in our results.

A complete taxonomic profile of all sample replicates at each individual time point is provided in Supplementary Information. An important aside to be noted with this data is the inherent reproducibility between sample triplicates. This finding provides a level of reassurance with regards to the aggregate nature of these water samples, often neglected in similar studies.

### Carbon cycling

Initial time points were characterised by a significantly (*p* ≤ 0.05) higher relative abundance of RXNs using molecular dioxygen as substrate such as: cytochrome-c oxidase (CYTOCHROME-C-OXIDASE-RXN, EC:1.9.3.1, Figure 5A), quinol–cytochrome-c reductase (1.10.2.2-RXN) and several mono- and dioxygenases. This was combined with a significantly (*p* ≤ 0.05) lower relative abundance of methanogenesis-related RXNs, nitrogenase (ferredoxin, NITROGENASE-RXN), rubisco (RIBULOSE-BISPHOSPHATECARBOXYLASE-RXN), or heterolactic fermentation (PHOSPHOKETOLASE-RXN). Catalase (CATAL-RXN, EC:1.11.1.6/21, see Reactive oxygen species detoxification in SI) gene relative abundance was highest at day 4 along with several ABC transporters and phosphotransferases, e.g. D-ribopyranose, D-xylose, spermidine or L-arginine, as well as assimilatory sulfur and nitrogen pathway RXNs (refer to SI for details). The higher relative abundances of all these genes at day 4 suggest a metabolism more dependent on oxygen, i.e. aerobic, or at least microaerophilic (see Nitrogen cycling). It also confers a higher dependence on available organic carbon (viz. heterotrophy), or on the presence of decaying complex organic matter (48). These interpretations are collectively supported by the geochemical measurements including the higher *E_h_* values, decreasing concentrations of sulfur and absence of nitrate, along with lower iron concentrations across the first sampling points.

**Figure 5.**
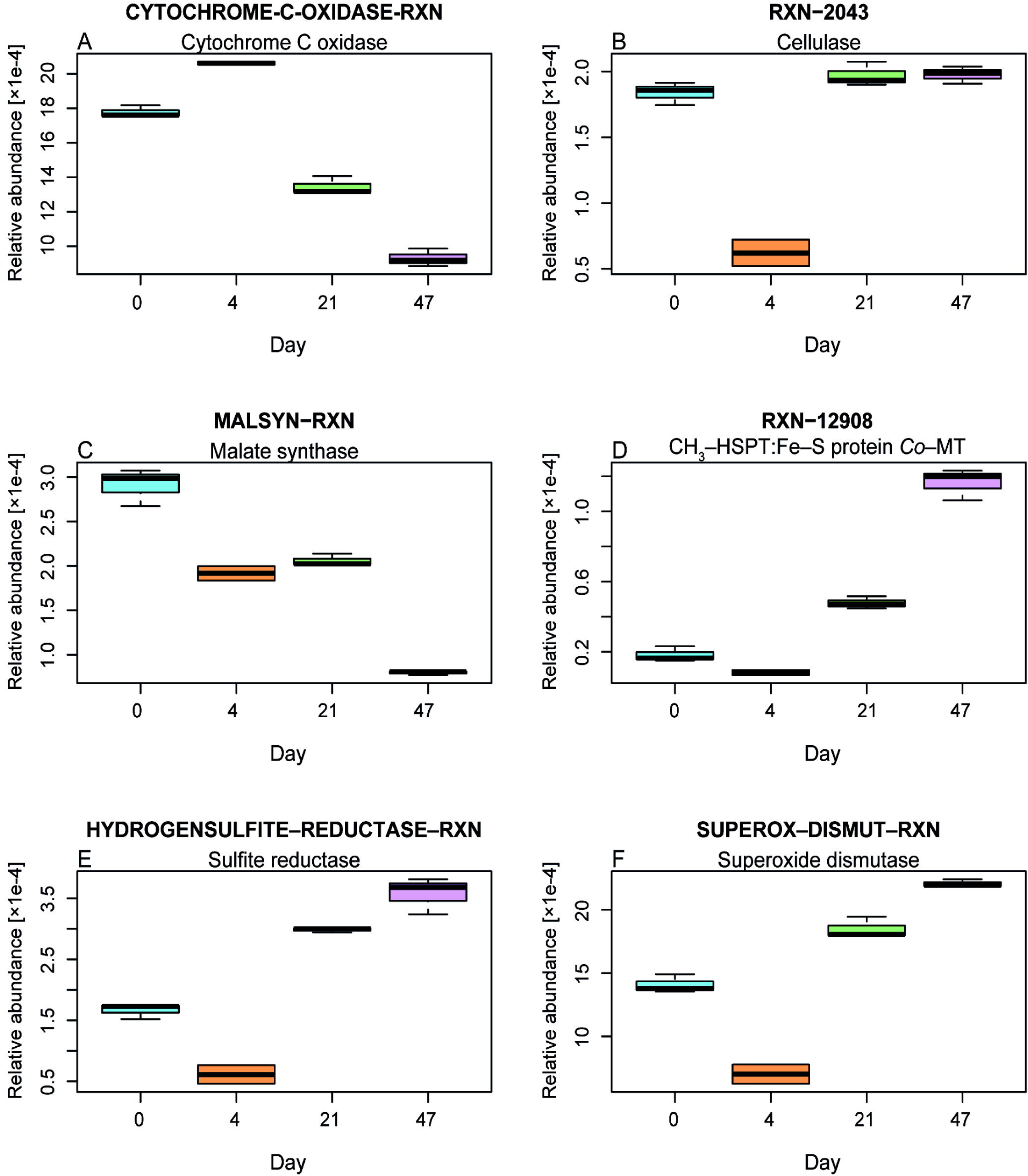
Changes in the relative abundances of selected RXNs over time. A, Cytochrome C oxidase; B, cellulase; C, malate synthase; D, 5-methyltetrahydrosarcinapterin:corrinoid/iron-sulfur protein Co-methyltransferase (CH3-HSPT:Fe-S protein Co-MT); E: sulfite reductase; and F, superoxide dismutase.

Regarding the source of the organic fraction, the results suggest that soil particles and associated organo-chemicals were mobilised from material above the trenches via advective transport mechanisms during and immediately after rainfall. This was evidenced by the increased proportion of soil-associated *Actinobacteria* at day 0, which diminished to 0.71% by day 4 (Figure 4). Furthermore, the presence of genes encoding for enzymes representing chitin degradation, such as chitobiase (RXN-12625, EC:3.2.1.52), diminished progressively until day 47 indicating a potential one-off provision of chitin, matching the leaching hypothesis (Figure S2). In addition, numerous facultative and strict anaerobes can degrade chitin and/or chitobiose (49–51), which lessens the likelihood of the alternative hypothesis that chitin depletion could be derived from the intrinsic lack of chitobiase in anaerobes.

The other primary biopolymer found in soil is cellulose (52). The relative abundance of cellulase (RXN-2043, EC:3.2.1.4) showed no significant differences between days 0, 21 and 47 (*p* = 0.172) suggesting a consistent supply of cellulose (Figure 5B). The exception, noted at day 4, can be explained by the proliferation of *Burkholderiales*. The constant abundance of cellulose is unsurprising given the historical records which indicate a range of organic (particularly cellulose-based) compounds were co-disposed of in the LFLS trenches (11). This hypothesis accords with previous stable isotope measurements, which identified the degradation of legacy organic matter as being responsible for enriched δ^13^C (inorganic) values measured near the trench (11).

Lignin is present to a lesser or greater extent in all plant biomass. From the list of enzymes in the literature capable of degrading lignin (53, 54), only below-threshold levels of certain genes were detected: versatile peroxidase (RXN0-267, EC:1.11.1.16), dye peroxidase (RXN-11813, EC:1.11.1.19) and laccase (LACCASE-RXN, EC:1.10.3.2). The overall lack of genes for proteins capable of lignin degradation strengthens the hypothesis that trench waste material is the source of cellulosic material rather than being derived from the degradation of plant material from the topsoil.

The two fundamental genes representing the glyoxylate pathway, isocitrate lyase (ISOCIT-CLEAV-RXN, EC:4.1.3.1, Figure S3) and malate synthase (MALSYN-RXN, EC:4.1.3.2, Figure 5C) showed between 3–4 times higher relative abundance in days 0 and 4 when compared to day 47. This pathway facilitates the use of short chain carbon compounds (C2–C3) for anabolic purposes, compounds especially generated during anaerobic processes. Although the glyoxylate pathway genes have been previously detected in organisms living in anaerobic environments (55), it was absent from strict anaerobic organisms based on the results from EggNOG (56), KEGG (57) and UniProtKB (58) databases (as of 24/05/2017). This finding supports the hypothesis that trench waters became anoxic as the water level declined, promoting conditions which favour the development of strict anaerobes. Furthermore, the aerobic community which developed immediately after the rainfall event likely utilised (and benefited from) the short chain carbon compounds generated by preceding anaerobic conditions.

Both acetoclastic and hydrogenotrophic methanogenesis pathways were evident from the functional profile analysis. Similar maximum relative abundances, about 1.4·10^-4^, of 5-methyltetrahydrosarcinapterin:corrinoid/iron-sulfur protein *Co*-methyltransferase (RXN-12908, EC:2.1.1.245, Figure 5D) and coenzyme F420-dependent hydrogenase (COENZYME-F420-HYDROGENASE-RXN, EC:1.12.98.1, Figure S4), which are specific RXNs for the methanogenesis from acetate and H_2_ and CO_2_, were observed at day 47. The presence of methanogenesis-specific RXNs is supported by previous isotopic measurements, showing δ^2^H enrichment, relative to δ^18^O, in the vicinity of the trenches (11). Concurrently, the taxonomy revealed the presence of the ANME-2d division contributing up to 13.8% of all *Archaea* at day 47 (1.5% of the community). The ANME-2d division was initially linked to the anaerobic oxidation of methane using NO_3_^-^ (42) or even NO_2_^-^ (59), constituting one of the primary methane sinks in anaerobic conditions. However, recent research has shown that some of the members of this taxa are able to utilise Fe(III) in place of NO_2_^-^ or NO_3_^-^ (44, 60, 61). This provides a more reasonable explanation given the limited nitrate present and abundance of iron during the later sampling days.

### Sulfur cycling

The relative abundance of sulfite cytochrome *c* reductase (SULFITEDEHYDROGENASE-RXN, EC:1.8.2.1, Figure S5), involved in the assimilatory reduction of sulfate, peaked at day 4, with ~10 times higher relative abundance compared to day 0. At the same time, dissimilatory sulfate reduction RXNs, e.g. dissimilatory sulfite reductase (HYDROGENSULFITE-REDUCTASE-RXN, EC:1.8.99.3), followed the opposite trend, with a maximum at day 47 (Figure 5E). The combined relative abundance of all the putative sulfate reducing bacteria (SRB) taxa detected (mainly *Syntrophobacterales*, *Thermodesulfovibrionaceae* and *Desulfobacterales*) also increased over time, from 2.3% of the total prokaryotes at day 0 to 4.9% at day 47. Dissimilatory sulfate reduction is one of the major redox processes in both natural (62) and artificial (63) anaerobic environments. The ten-fold reduction in the sulfur concentration is thought to result from the loss of dissolved sulfate via reduction to sulfide and subsequent precipitation of insoluble heavy metal sulfides, particularly FeS. This interpretation is supported by the lack of measurable sulfides in the trench water (see Chemical analyses of trench waters), along with previous reports showing sulfate reduction based on isotopic fractionation and severe (10 to 100-fold) depletion in sulfate concentrations in trench waters relative to surrounding wells (11). The potential contribution of sulfate- and nitrite-dependent anaerobic methane oxidation (S-DAMO and N-DAMO respectively) was discounted due to the near complete absence of ANME taxa (*Archaea*) aside from ANME-2d, and NC10 (*Bacteria*) (see Nitrogen cycling) (44, 59, 64, 65).

### Nitrogen cycling

The (inorganic) nitrogen cycle was mostly represented by RXNs related to denitrification and assimilatory nitrogen reactions, primarily dissimilatory nitrate reductase (RXN-16471, EC:1.7.5.1, see Figure S6 for a complete profile of the N-associated RXNs), although nitrogen fixation (NITROGENASE-RXN, EC:1.18.6.1) was the predominant N-associated RXN. All other prominent N-associated RXNs peaked at day 4 with the exception of hydroxylamine reductase (HYDROXYLAMINE-REDUCTASE-RXN, EC:1.7.99.1) and ammonia oxygenase (AMONITRO-RXN, EC:1.14.99.39). The increase in nitrate concentrations observed between days 4 and 6 (0.236 and 0.552 μM respectively) might be derived from the oxidation of NO by nitric oxide dioxygenase (R621-RXN, EC:1.14.12.17). This surge in the relative abundance of nitrogen metabolism RXNs (at day 4) coincided with an increased dominance of *Betaproteobacteria*, mostly *Burkholderiales*. *Betaproteobacteria* have been linked to denitrification processes in a uranium-polluted aquifer in Rifle (USA) (66) and they may play a similar ecological role at LFLS. However, the increase in nitrate concentrations could equally be indicative of NO or NO_2_^-^ oxidation, originally produced by the aerobic oxidation of ammonium by the *Thaumarchaeota* (67, 68).

The inorganic nitrogen cycling results provide important information regarding the transition from aerobic to anaerobic conditions as the water level declines. The competing requirements of nitrate respiration, which necessitates oxygen-limiting conditions at a minimum (69), and NO dioxygenase needing molecular oxygen, collectively suggest that at day 4 the trench waters were no longer aerobic, but more likely microaerophilic/hypoxic. However, many facultative anaerobes are capable of using nitrate as an alternate electron acceptor when oxygen is not available. As such, the high relative abundance of nitrate reducing enzyme genes at day 4 (viz. dissimilatory nitrate reductases) could be a confounding effect associated with the increased abundance of *Burkholderiales*. This was confirmed by searching the denitrifying pathway in KEGG (map00910) (as of 08/07/2016) (57).

### Iron cycling

Temporal increases in Fe(II) concentrations, such those experienced in the later anoxic periods of the sampling, may be associated with the use of Fe(III) during anaerobic respiration, previously attributed to decaheme *c*-type cytochromes from the OmcA/MtrC family present in iron-reducing bacteria (FeRB) such as *Shewanella* spp. and *Geobacter* spp. (70). However, their representation in the trench water metagenome data was scarce (relative abundance ≤ 10^-6^). It is known that for respiration to occur, FeRB often require direct physical contact with Fe(III) solids (71). As only aqueous/suspended phase sampling was permissible at LFLS, it is possible the under representation of a FeRB community was due to our sampling regime in which the solid-phase was largely excluded.

The genes associated with RXNs specific to the Fe(II) oxidation pathway (via rusticyanin, PWY-6692): ubiquinone-cytochrome-*bc1* reductase (EC:1.10.2.2, RXN-15829) and *aa3*-type cytochrome oxidase (EC:1.9.3.1, RXN-15830) were present, despite no ironrusticyanin reductase (EC:1.16.9.1, RXN-12075), *cyc2*, being found under the established threshold (maximum value ~3×10^-6^) (72). The presence of both cytochromes could be because HUMAnN2 could not distinguish between the ‘generic’ quinol–cytochrome-c reductase (RXN-15816, EC:1.10.2.2), and cytochrome *c* oxidase (CYTOCHROME-COXIDASE-RXN) respectively, particularly when analysing short reads.

The presence of 1.9% of *Gallionellaceae* (over the total community composition) at day 0 and 2.9% of *Crenothrix*, a non-FeOB usually associated with Fe-mineralising biofilms (73), suggest the existence of transient microaerophilic iron oxidising activity in the trench water despite the extremely low relative abundance of *cyc2*. Waste disposal records from LFLS indicate that numerous steel drums (~760) were co-disposed of in the trenches (36). Based on the active sulfate respiration observed in the trench waters, we would expect that iron/steel materials may have suffered microbially induced corrosion (MIC) (74), potentially contributing to the elevated Fe(II) concentrations. However, the main iron MIC product is likely FeS (74), which can be utilised by *Gallionella ferruginea* (*Gallionellaceae*), an iron oxidising bacterium (FeOB) (75). Furthermore, this process would also provide an explanation for a persistent source of Fe(III) (oxy)hydroxides and their ongoing interaction with the S-cycle.

Previous research on soils experiencing fluctuating redox conditions and active iron cycling have shown they frequently lack ‘typical’ Fe(III)-respiring bacteria (76), being outcompeted by sulfate respiration (77). Despite Fe(III) being a more thermodynamically favourable electron acceptor it is commonly acknowledged that sulfate respiration often outcompetes Fe(III) respiration in both high and low sulfate environments (77, 78). High levels of sulfate reduction in low-sulfate environments, akin to the LFLS trenches, has been previously observed to occur in the presence of crystalline Fe(III) oxyhydroxides that partially re-oxidise sulfide generated by SRB to elemental sulfur (78). This mechanism provides a plausible explanation for our observations at LFLS. The large concentration of dissolved Fe(II) in the trench waters would likely contribute to the Fe(II)-catalysed transformation of amorphous ferrihydrite to more crystalline, thermodynamically-favourable forms (79, 80), although this would be limited to some extent by the large concentrations of dissolved organic matter and silica (81). Furthermore, the abiotic reduction of crystalline Fe(III) oxyhydroxides would severely limit the energy acquisition by FeRB and therefore the FeRB would only outcompete SRB when sulfate and other reducible sulfur compounds were totally consumed.

Alternatively, fermentative bacteria coupling the oxidation of a range of organic compounds to the reduction of ferric iron (82–84) may play an integral role in maintaining the elevated Fe(II) concentrations observed within the trenches. One example is the organism *Propionibacterium freudenreichii* which has previously been observed to reduce Fe(III) by using humic substances as an electron mediator (85). In this regard, both heterolactic and propionic acid fermentation pathways showed high relative abundances based on phosphoketolase (PHOSPHOKETOLASE-RXN, EC:4.1.2.9) and methylmalonyl-CoA decarboxylase (RXN0-310, EC:4.1.1.41), respectively (Figure S7). Their decrease in relative abundance (at day 4) could be explained by the lack of fermentative pathways present in the genomes associated with the population shift, i.e. increase in *Burkholderiales*. This was confirmed by searching (as of 08/06/2016) methylmalonyl-CoA decarboxylase and phosphoketolase in the KEGG maps (map00640 and map00030, respectively).

### Synthesis of trench processes and radiochemical mobilisation implications

The biogeochemistry of this dynamic system is conceptually described by the elemental cycling schematic shown in Figure 6. Immediately following the rainfall event, the *E_h_* reached its most oxidising value (247 mV) with the microbial community characterised by higher relative abundances of pathways related to aerobic (or at least microaerophilic) heterotrophic metabolism and one capable of chitin degradation. Although conditions were not strongly oxidising, this pulse of oxic water would be sufficient to induce the abiotic oxidation of part of the large store of dissolved Fe(II), likely forming a combination of the reactive Fe(III) oxyhydroxides ferrihydrite and silica-ferrihydrite, along with the more crystalline oxyhydroxide lepidocrocite, as has previously been observed in these trench waters (37). As both Am(III) and Pu(III)/(IV) are known to strongly sorb to sediments and iron oxides (15), it is of little surprise to observe the greatest proportions of (suspended) solid-associated actinides at this time point (day 0, Figure 3). Note that even though most Am (54.3%) and particularly Pu (78.8%) were associated with a solid-fraction >0.45 μm, they were still extracted from the trench under our low-flow sampling method. This finding implies that colloid-associated Pu, and to a lesser extent Am, remains mobile within the trench waters, congruent with observations from other legacy radioactive waste sites (86). Interestingly though, the Pu migration distance away from source at LFLS in groundwater has been shown to be much smaller (~1 order of magnitude less) compared to other legacy locations such as the Nevada Test Site (87), Rocky Flats (88) and Mayak (89). We attribute this to a low permeability soil matrix, inhibiting downward migration to the connected/permanent water table, coupled with the biogeochemical conditions within the trenches themselves. The high concentrations of Fe(II), circumneutral pH and proliferation of aerobic heterotrophs drives the rapid formation of large quantities of Fe(III) oxides. The quantity of Fe(III) oxides that form upon oxic rainwater intrusion is evidently sufficient to contain the bulk of contaminants within the trenches during ‘bathtub’ overflow events. The rate of Fe(II) oxidation is likely to be crucial for the ongoing attenuation of Pu and Am at LFLS (37).

**Figure 6.**
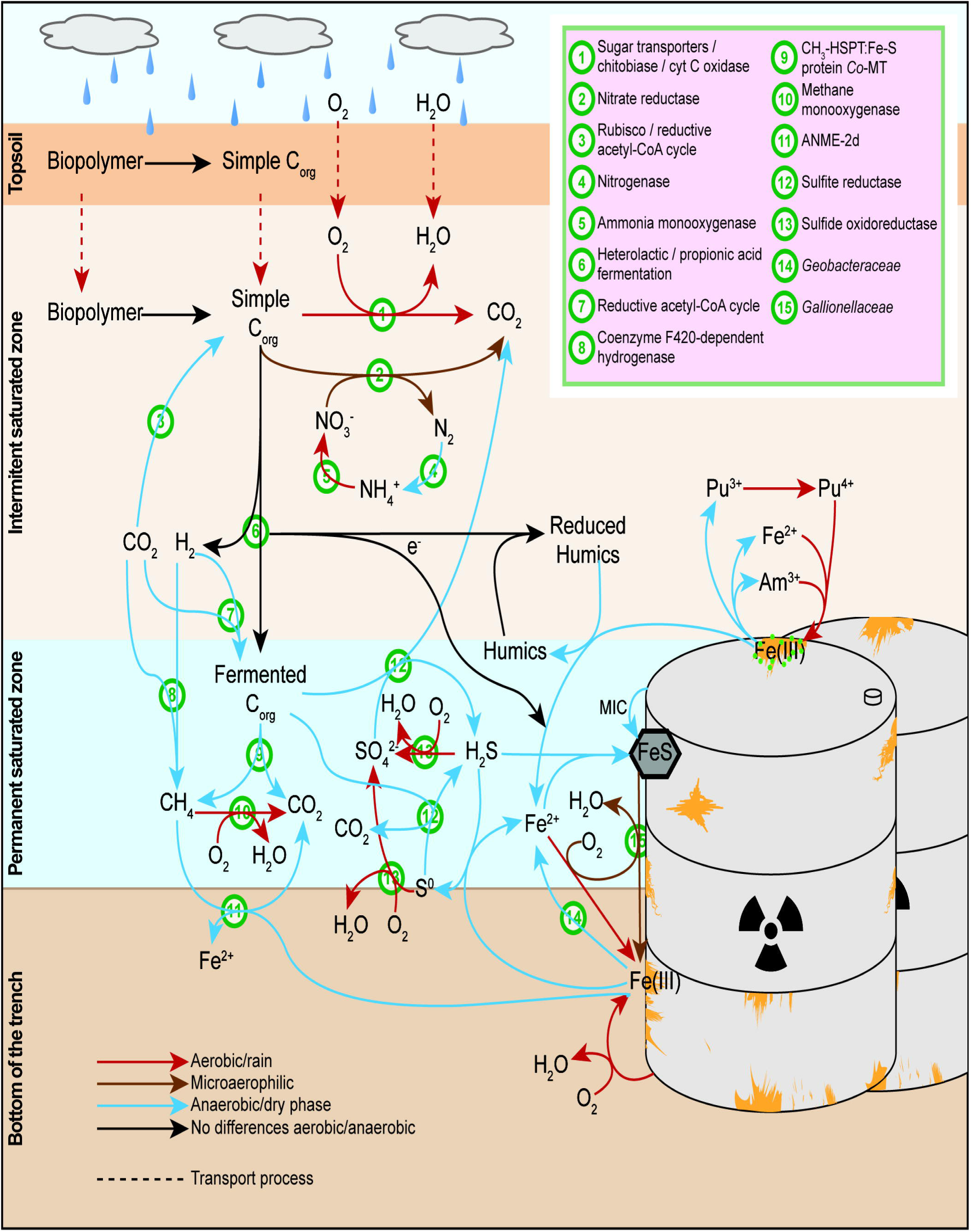
Hypothetical global scheme of the processes at LFLS. Orange details on barrels reading Fe(III) represent solid Fe(III) minerals, mainly (oxy)hydroxides. Colour of arrows represent the time at which each process takes place: red, aerobic/after rain; brown, microaerophilic; blue, anaerobic/dry phase; and black arrows indicate processes that seem to be independent from the sampling time. Dashed arrows indicate transport. Abbreviations used in the figure: C_org_, organic carbon; MIC, microbially induced corrosion; CH_3_-HSPT:Fe-S protein *Co*-MT, 5-methyltetrahydrosarcinapterin:corrinoid/iron-sulfur protein Co-methyltransferase.

By day 4, as water levels rapidly decrease, an increase of *Burkholderiales* (*Betaproteobacteria*) generates a functional profile disturbance and, while the aerobic profile is maintained, it is also the time-point at which nitrogen cycling (e.g. nitrate respiration) becomes most active (Figure 6; ⍰). Therefore, day 4 represents a potential transition away from oxic conditions derived from rain infiltration, though this sequence of events is somewhat confounded by the microbial growth lag phase associated with aerobic respirators.

Over the following weeks (days 21 and 47), the microbial community transitions to a functional profile dominated by carbon fixation, methanogenesis and sulfate respiration pathways (Figure 6; ⍰⍰⍰⍰). The increase in anaerobicity correlated with increasing concentrations of soluble Fe and soluble radionuclides and a depletion of sulfate and nitrate. In the case of Am, although the total activity increased gradually as the water level in the trench declined, the soluble fraction increased by a greater proportion (Figure 3). This indicates that either a desorption or dissolution process occurred. The concomitant increase in the soluble Am fraction and Fe(II) concentrations point towards reductive dissolution of Fe(III) oxyhydroxides as a major driver behind Am solubilisation.

The Pu behaviour in the trenches presents a more complex temporal dynamic due, in part, to its more varied redox chemistry. As conditions become more reducing over time, one would expect to observe soluble Pu activities increase, through both the dissolution of Fe(III) oxides and, to a lesser extent, from Pu(IV) to Pu(III) reduction. In an interesting point of difference to Am, our results show that Pu proportionally remained in the particle-associated fraction for a substantially longer period, even as the dissolved Am(III) activity and Fe(II) concentration increased. The reason for the differing solution/solid-phase partitioning observed for Pu and Am is not clear based on the evidence to hand. The inability to sample solid materials from within the trenches has limited more conclusive understanding. However, we suggest the differing Am and Pu associations are likely due to different redox states, based on the previous measurements at LFLS, which showed a dominant Pu oxidation state of +4 (29). Electrostatically, Pu(IV), as the neutral hydrolysed cation Pu(OH)_4_(aq) or as an intrinsic colloid, is more likely sorbed or incorporated into positively charged pseudo-colloids such as Fe(III) oxides, compared to Am^3+^. This is supported by research showing the highly reversible nature of Am(III) sorption onto poor crystalline iron colloids (90) and association with carbonate- and exchangeable sites on clays (91).

## Conclusions

The inability to comprehensively access and sample within legacy radioactive waste environments hampers our ability to comprehend co-occurring elemental cycling and microbial metabolism, potentially curtailing our ability to effectively manage and remediate such sites. The trench-sampling point at LFLS is therefore a particularly useful resource for such research. In this study, our coupled use of metagenomics and chemical analyses has provided a previously unattainable level of understanding for the LFLS trench water, highlighting the responsiveness of the microbial community to external changes and dynamic nature of the resulting chemistry. The combined results show that the trench waters contain a taxonomically diverse microbial community, which has likely evolved in response to variations in energy sources supplied by frequent redox fluctuations. When combined with the complex nature of the waste-form, a myriad of micro-environments has developed within the trenches, allowing for simultaneous O, N, Fe, S and C elemental cycling, as shown by co-occurring metabolic reactions in the aggregate water samples. Consequently, it can be inferred that Pu and Am are subject to persistent reducing conditions (as evident from active iron oxide dissolution, sulfate reduction and methanogenesis) when the water level is low between rainfall events. These reductive processes maintain Pu and Am solubility, despite the occasional onset of oxidising conditions associated with rainfall events. Ultimately however, the high concentrations of Fe present and the tendency of Fe(II) to be relatively rapidly oxidised to strongly sorbing Fe(III) (oxy)hydroxide solids on exposure to oxic conditions results in limited transport of Pu and Am.

Although the findings described above are intrinsically linked to the specific site under investigation, they provide important generic insights into the dynamic biogeochemical behaviour of iron-rich, redox-cycling environments. Of particular interest is the rapid response of the microbial community to dynamic redox conditions and the potential impact upon persistent contaminant solubility and enhanced mobility.

## Acknowledgements

ANSTO staff members are thanked for their valuable contributions: Kerry Wilsher (fieldwork, laboratory analyses), Sangeeth Thiruvoth (actinides), Brett Rowling (fieldwork, inorganics), Henri Wong (inorganics), Chris Vardanega (inorganics) and Dioni Cendón (discussion). The Australian Bureau of Meteorology is acknowledged for rainfall data.

## Author contribution

Study concept and experimental design: XVC, ASK, TDW. Fieldwork: ASK. Chemical and radiochemical analyses: ASK, JJH. Molecular biology and bioinformatics analyses: XVC. Contribution of reagents/materials/analysis tools: TEP, TDW. Manuscript writing: XVC, ASK. Revision and review of the manuscript: XVC, ASK, MWB, TDW.

## References

1. Cleveland JM, Rees TF. 1981. Characterization of plutonium in Maxey Flats radioactive trench leachates. Science 212:1506–1509.

2. Saunders JA, Toran LE. 1995. Modeling of radionuclide and heavy metal sorption around low- and high-pH waste disposal sites at Oak Ridge, Tennessee. Appl Geochem 10:673–684.

3. Cantrell KJ, Felmy AR. 2012. Plutonium and americium geochemistry at Hanford: a site-wide review. RPT-DVZ-AFRI-001 PNNL-21651. Pacific Northwest National Laboratory, Washington (USA).

4. Gerber MS. 1992. Past practices technical characterization study — 300 area — Hanford site. WHC-MR-0388. UC-702. Westinghouse Hanford Company, Richland, WA.

5. Killey RWD, Rao RR, Eyvindson S. 1998. Radiocarbon speciation and distribution in an aquifer plume and groundwater discharge area, Chalk River, Ontario. Appl Geochem 13:3–16.

6. Fellingham L, Graham A, Stiff S. 2003. Characterisation and remediation of beryllium waste pits in the Southern storage area at Harwell, p. 1041–1049. In ASME 2003 9th International Conference on Radioactive Waste Management and Environmental Remediation. American Society of Mechanical Engineers, Oxford, England.

7. Wilkins MJ, Livens FR, Vaughan DJ, Lloyd JR, Beadle I, Small JS. 2010. Fe(III) reduction in the subsurface at a low-level radioactive waste disposal site. Geomicrobiol J 27:231–239.

8. Gudelis A, Gvozdait R, Kubarevi ien V, Druteikien R, Lukoševi ius Š, Šutas A. 2010. On radiocarbon and plutonium leakage to groundwater in the vicinity of a shallow-land radioactive waste repository. J Environ Radioact 101:443–445.

9. Bugai D, Kashparov V, Dewiére L, Khomutinin Y, Levchuk S, Yoschenko V. 2005. Characterization of subsurface geometry and radioactivity distribution in the trench containing Chernobyl clean-up wastes. Environ Geol 47:869–881.

10. Payne TE, Harrison JJ, Hughes CE, Johansen MP, Thiruvoth S, Wilsher KL, Cendón DI, Hankin SI, Rowling B, Zawadzki A. 2013. Trench ‘bathtubbing’ and surface plutonium contamination at a legacy radioactive waste site. Environ Sci Technol 47:13284–13293.

11. Cendón DI, Hughes CE, Harrison JJ, Hankin SI, Johansen MP, Payne TE, Wong H, Rowling B, Vine M, Wilsher K, Guinea A, Thiruvoth S. 2015. Identification of sources and processes in a low-level radioactive waste site adjacent to landfills: groundwater hydrogeochemistry and isotopes. Aust J Earth Sci 62:123–141.

12. Hughes CE, Cendón DI, Harrison JJ, Hankin SI, Johansen MP, Payne TE, Vine M, Collins RN, Hoffmann EL, Loosz T. 2011. Movement of a tritium plume in shallow groundwater at a legacy low-level radioactive waste disposal site in eastern Australia. J Environ Radioact 102:943–952.

13. Isaacs SR, Mears KF. 1977. A study of the burial ground used for radioactive waste at the Little Forest area near Lucas Heights New South Wales. AAEC/E--427. Australian Atomic Energy Commission Research Establishment, Lucas Heights, N.S.W. (AU).

14. Pett-Ridge J, Firestone MK. 2005. Redox fluctuation structures microbial communities in a wet tropical soil. Appl Environ Microbiol 71:6998–7007.

15. Choppin GR. 2007. Actinide speciation in the environment. J Radioanal Nucl Chem 273:695–703.

16. Kimber RL, Boothman C, Purdie P, Livens FR, Lloyd JR. 2012. Biogeochemical behaviour of plutonium during anoxic biostimulation of contaminated sediments. Mineral Mag 76:567–578.

17. Lloyd JR, Sole VA, Praagh CVGV, Lovley DR. 2000. Direct and Fe(II)-mediated reduction of technetium by Fe(III)-reducing bacteria. Appl Environ Microbiol 66:3743–3749.

18. Renshaw JC, Butchins LJC, Livens FR, May I, Charnock JM, Lloyd JR. 2005. Bioreduction of uranium: Environmental implications of a pentavalent intermediate. Environ Sci Technol 39:5657–5660.

19. Francis AJ, Dodge CJ, Gillow JB. 2008. Reductive dissolution of Pu(IV) by *Clostridium* sp. under anaerobic conditions. Environ Sci Technol 42:2355–2360.

20. Renshaw JC, Law N, Geissler A, Livens FR, Lloyd JR. 2009. Impact of the Fe(III)-reducing bacteria *Geobacter sulfurreducens* and *Shewanella oneidensis* on the speciation of plutonium. Biogeochemistry 94:191–196.

21. Fomina M, Charnock JM, Hillier S, Alvarez R, Gadd GM. 2007. Fungal transformations of uranium oxides. Environ Microbiol 9:1696–1710.

22. Wilkins MJ, Livens FR, Vaughan DJ, Beadle I, Lloyd JR. 2007. The influence of microbial redox cycling on radionuclide mobility in the subsurface at a low level radioactive waste storage site. Geobiology 5:293–301.

23. Vázquez-Campos X, Kinsela AS, Collins RN, Neilan BA, Aoyagi N, Waite TD. 2015. Uranium binding mechanisms of the acid-tolerant fungus *Coniochaeta fodinicola*. Environ Sci Technol 49:8487–8496.

24. Amachi S, Fujii T, Shinoyama H, Muramatsu Y. 2005. Microbial influences on the mobility and transformation of radioactive iodine in the environment. J Nucl Radiochem Sci 6:21–24.

25. Bertin PN, Heinrich-Salmeron A, Pelletier E, Goulhen-Chollet F, Arsène-Ploetze F, Gallien S, Lauga B, Casiot C, Calteau A, Vallenet D, Bonnefoy V, Bruneel O, Chane-Woon-Ming B, Cleiss-Arnold J, Duran R, Elbaz-Poulichet F, Fonknechten N, Giloteaux L, Halter D, Koechler S, Marchal M, Mornico D, Schaeffer C, Smith AAT, Van Dorsselaer A, Weissenbach J, Médigue C, Le Paslier D. 2011. Metabolic diversity among main microorganisms inside an arsenic-rich ecosystem revealed by meta- and proteogenomics. ISME J 5:1735–1747.

26. Castelle CJ, Hug LA, Wrighton KC, Thomas BC, Williams KH, Wu D, Tringe SG, Singer SW, Eisen JA, Banfield JF. 2013. Extraordinary phylogenetic diversity and metabolic versatility in aquifer sediment. Nat Commun 4:2120.

27. Aguiar-Pulido V, Huang W, Suarez-Ulloa V, Cickovski T, Mathee K, Narasimhan G. 2016. Metagenomics, metatranscriptomics, and metabolomics approaches for microbiome analysis. Evol Bioinform 12:5–16.

28. Allen EE, Banfield JF. 2005. Community genomics in microbial ecology and evolution. Nat Rev Micro 3:489–498.

29. Ikeda-Ohno A, Harrison JJ, Thiruvoth S, Wilsher K, Wong HKY, Johansen MP, Waite TD, Payne TE. 2014. Solution speciation of plutonium and americium at an Australian legacy radioactive waste disposal site. Environ Sci Technol 48:10045–10053.

30. Harrison JJ, Zawadzki A, Chisari R, Wong HKY. 2011. Separation and measurement of thorium, plutonium, americium, uranium and strontium in environmental matrices. J Environ Radioact 102:896–900.

31. APHA, AWWA, WEF. 1998. Standard methods for the examination of water and wastewater20th ed. APHA-AWWA-WEF, Washington, D.C.

32. Jacobs J, Rhodes M, Sturgis B, Wood B. 2009. Influence of environmental gradients on the abundance and distribution of *Mycobacterium* spp. in a coastal lagoon estuary. Appl Environ Microbiol 75:7378–7384.

33. Abubucker S, Segata N, Goll J, Schubert AM, Izard J, Cantarel BL, Rodriguez-Mueller B, Zucker J, Thiagarajan M, Henrissat B, White O, Kelley ST, Methé B, Schloss PD, Gevers D, Mitreva M, Huttenhower C. 2012. Metabolic reconstruction for metagenomic data and its application to the human microbiome. PLoS Comput Biol 8:e1002358.

34. Parks DH, Tyson GW, Hugenholtz P, Beiko RG. 2014. STAMP: statistical analysis of taxonomic and functional profiles. Bioinformatics 30:3123–3124.

35. Benjamini Y, Hochberg Y. 1995. Controlling the False Discovery Rate: A Practical and Powerful Approach to Multiple Testing. J Roy Stat Soc B 57:289–300.

36. Payne TE. 2012. Background report on the Little Forest Burial Ground legacy waste site. ANSTO / E-780. Institute for Environmental Research, Australian Nuclear Science and Technology Organisation, Lucas Heights, N.S.W. (AU).

37. Kinsela AS, Jones AJ, Bligh MW, Pham AN, Collins RN, Harrison JJ, Wilsher KL, Payne TE, Waite TD. 2016. Influence of dissolved silicate on Fe(II) oxidation. Environ Sci Technol 50:11663–11671.

38. Jiang L, Zheng Y, Chen J, Xiao X, Wang F. 2011. Stratification of archaeal communities in shallow sediments of the Pearl River Estuary, Southern China. Antonie van Leeuwenhoek 99:739–751.

39. Meng J, Xu J, Qin D, He Y, Xiao X, Wang F. 2014. Genetic and functional properties of uncultivated MCG archaea assessed by metagenome and gene expression analyses. ISME J 8:650–659.

40. Evans PN, Parks DH, Chadwick GL, Robbins SJ, Orphan VJ, Golding SD, Tyson GW. 2015. Methane metabolism in the archaeal phylum Bathyarchaeota revealed by genome-centric metagenomics. Science 350:434–438.

41. He Y, Li M, Perumal V, Feng X, Fang J, Xie J, Sievert SM, Wang F. 2016. Genomic and enzymatic evidence for acetogenesis among multiple lineages of the archaeal phylum Bathyarchaeota widespread in marine sediments. Nat Microbiol 16035.

42. Haroon MF, Hu S, Shi Y, Imelfort M, Keller J, Hugenholtz P, Yuan Z, Tyson GW. 2013. Anaerobic oxidation of methane coupled to nitrate reduction in a novel archaeal lineage. Nature 500:567–570.

43. Karakashev D, Batstone DJ, Trably E, Angelidaki I. 2006. Acetate oxidation is the dominant methanogenic pathway from acetate in the absence of methanosaetaceae. Appl Environ Microbiol 72:5138–5141.

44. Leu A, Hu S, Cai C, Yuan Z, Orphan VJ, Tyson GW. 2016. Understanding the metabolic potential of ANME-2d and its role in anaerobic methane oxidation coupled to metal reduction. Montreal, Canada.

45. Luef B, Frischkorn KR, Wrighton KC, Holman H-YN, Birarda G, Thomas BC, Singh A, Williams KH, Siegerist CE, Tringe SG, Downing KH, Comolli LR, Banfield JF. 2015. Diverse uncultivated ultra-small bacterial cells in groundwater. Nat Commun 6:6372.

46. Baker BJ, Tyson GW, Webb RI, Flanagan J, Hugenholtz P, Allen EE, Banfield JF. 2006. Lineages of acidophilic Archaea revealed by community genomic analysis. Science 314:1933–1935.

47. Brown CT, Hug LA, Thomas BC, Sharon I, Castelle CJ, Singh A, Wilkins MJ, Wrighton KC, Williams KH, Banfield JF. 2015. Unusual biology across a group comprising more than 15% of domain Bacteria. Nature 523:208–211.

48. Peng R-H, Xiong A-S, Xue Y, Fu X-Y, Gao F, Zhao W, Tian Y-S, Yao Q-H. 2008. Microbial biodegradation of polyaromatic hydrocarbons. FEMS Microbiol Rev 32:927–955.

49. Riemann L, Azam F. 2002. Widespread *N*-acetyl-D-glucosamine uptake among pelagic marine bacteria and its ecological implications. Appl Environ Microbiol 68:5554–5562.

50. Sorokin DY, Rakitin AL, Gumerov VM, Beletsky AV, Damsté S, S J, Mardanov AV, Ravin NV. 2016. Phenotypic and genomic properties of *Chitinispirillum alkaliphilum* gen. nov., sp. nov., a haloalkaliphilic anaerobic chitinolytic bacterium representing a novel class in the phylum *Fibrobacteres*. Front Microbiol 7.

51. Zavarzina DG, Tourova TP, Kolganova TV, Boulygina ES, Zhilina TN. 2009. Description of *Anaerobacillus alkalilacustre* gen. nov., sp. nov.—Strictly anaerobic diazotrophic bacillus isolated from soda lake and transfer of *Bacillus arseniciselenatis*, *Bacillus macyae*, and *Bacillus alkalidiazotrophicus* to *Anaerobacillus* as the new combinations *A. arseniciselenatis* comb. nov., *A. macyae* comb. nov., and *A. alkalidiazotrophicus* comb. nov. Microbiology 78:723–731.

52. Beier S, Bertilsson S. 2013. Bacterial chitin degradation—mechanisms and ecophysiological strategies. Front Microbiol 4:149.

53. de Gonzalo G, Colpa DI, Habib MHM, Fraaije MW. 2016. Bacterial enzymes involved in lignin degradation. J Biotechnol 236:110–119.

54. Pollegioni L, Tonin F, Rosini E. 2015. Lignin-degrading enzymes. FEBS J 282:1190–1213.

55. Burow LC, Mabbett AN, Blackall LL. 2008. Anaerobic glyoxylate cycle activity during simultaneous utilization of glycogen and acetate in uncultured *Accumulibacter* enriched in enhanced biological phosphorus removal communities. ISME J 2:1040–1051.

56. Huerta-Cepas J, Szklarczyk D, Forslund K, Cook H, Heller D, Walter MC, Rattei T, Mende DR, Sunagawa S, Kuhn M, Jensen LJ, von Mering C, Bork P. 2016. eggNOG 4.5: a hierarchical orthology framework with improved functional annotations for eukaryotic, prokaryotic and viral sequences. Nucl Acids Res 44:D286–D293.

57. Kanehisa M, Sato Y, Kawashima M, Furumichi M, Tanabe M. 2016. KEGG as a reference resource for gene and protein annotation. Nucl Acids Res 44:D457–D462.

58. Magrane M, Uniprot Consortium. 2011. UniProt Knowledgebase: a hub of integrated protein data. Database 2011:bar009.

59. Cui M, Ma A, Qi H, Zhuang X, Zhuang G. 2015. Anaerobic oxidation of methane: an “active” microbial process. MicrobiologyOpen 4:1–11.

60. Timmers PHA, Welte CU, Koehorst JJ, Plugge CM, Jetten MSM, Stams AJM. 2017. Reverse methanogenesis and respiration in methanotrophic Archaea. Archaea 2017:e1654237.

61. Ettwig KF, Zhu B, Speth D, Keltjens JT, Jetten MSM, Kartal B. 2016. Archaea catalyze iron-dependent anaerobic oxidation of methane. PNAS 113:12792–12796.

62. Grein F, Ramos AR, Venceslau SS, Pereira IAC. 2013. Unifying concepts in anaerobic respiration: Insights from dissimilatory sulfur metabolism. Biochim Biophys Acta Bioenerg 1827:145–160.

63. Paulo LM, Stams AJM, Sousa DZ. 2015. Methanogens, sulphate and heavy metals: a complex system. Rev Environ Sci Biotechnol 14:537–553.

64. Scheller S, Yu H, Chadwick GL, McGlynn SE, Orphan VJ. 2016. Artificial electron acceptors decouple archaeal methane oxidation from sulfate reduction. Science 351:703–707.

65. He Z, Cai C, Wang J, Xu X, Zheng P, Jetten MSM, Hu B. 2016. A novel denitrifying methanotroph of the NC10 phylum and its microcolony. Sci Rep 6:32241.

66. Wrighton KC, Castelle CJ, Wilkins MJ, Hug LA, Sharon I, Thomas BC, Handley KM, Mullin SW, Nicora CD, Singh A, Lipton MS, Long PE, Williams KH, Banfield JF. 2014. Metabolic interdependencies between phylogenetically novel fermenters and respiratory organisms in an unconfined aquifer. ISME J 8:1452–1463.

67. Könneke M, Schubert DM, Brown PC, Hügler M, Standfest S, Schwander T, BorzyskowskiLS von, Erb TJ, Stahl DA, Berg IA. 2014. Ammonia-oxidizing archaea use the most energy-efficient aerobic pathway for CO2 fixation. PNAS 111:8239–8244.

68. Pester M, Schleper C, Wagner M. 2011. The Thaumarchaeota: an emerging view of their phylogeny and ecophysiology. Curr Opin Microbiol 14:300–306.

69. Fischer M, Falke D, Pawlik T, Sawers RG. 2014. Oxygen-dependent control of respiratory nitrate reduction in mycelium of *Streptomyces coelicolor* A3(2). J Bacteriol 196:4152–4162.

70. Lovley DR, Holmes DE, Nevin KP. 2004. Dissimilatory Fe(III) and Mn(IV) reduction. Adv Microb Physiol 49:219–286.

71. Weber KA, Achenbach LA, Coates JD. 2006. Microorganisms pumping iron: anaerobic microbial iron oxidation and reduction. Nat Rev Micro 4:752–764.

72. Kato S, Ohkuma M, Powell DH, Krepski ST, Oshima K, Hattori M, Shapiro N, Woyke T, Chan CS. 2015. Comparative genomic insights into ecophysiology of neutrophilic, microaerophilic iron oxidizing bacteria. Front Microbiol 6.

73. Emerson D, Fleming EJ, McBeth JM. 2010. Iron-oxidizing bacteria: an environmental and genomic perspective. Ann Rev Microbiol 64:561–583.

74. Anandkumar B, George RP, Maruthamuthu S, Parvathavarthini N, Mudali UK. 2016. Corrosion characteristics of sulfate-reducing bacteria (SRB) and the role of molecular biology in SRB studies: an overview. Corrosion Rev 34:41–63.

75. Lütters-Czekalla S. 1990. Lithoautotrophic growth of the iron bacterium *Gallionella ferruginea* with thiosulfate or sulfide as energy source. Arch Microbiol 154:417–421.

76. DeAngelis KM, Silver WL, Thompson AW, Firestone MK. 2010. Microbial communities acclimate to recurring changes in soil redox potential status. Environ Microbiol 12:3137–3149.

77. Cozzarelli IM, Weiss JV. 2007. Biogeochemistry of Aquifer Systems, p. 843–859. In Hurst, CJ, Crawford, RL, Garland, JL, Lipson, DA, Mills, AL, Stetzenbach, LD (eds.), Manual of environmental microbiology. American Society of Microbiology.

78. Hansel CM, Lentini CJ, Tang Y, Johnston DT, Wankel SD, Jardine PM. 2015. Dominance of sulfur-fueled iron oxide reduction in low-sulfate freshwater sediments. ISME J 9:2400–2412.

79. Boland DD, Collins RN, Miller CJ, Glover CJ, Waite TD. 2014. Effect of solution and solid-phase conditions on the Fe(II)-accelerated transformation of ferrihydrite to lepidocrocite and goethite. Environ Sci Technol 48:5477–5485.

80. Pedersen HD, Postma D, Jakobsen R, Larsen O. 2005. Fast transformation of iron oxyhydroxides by the catalytic action of aqueous Fe(II). Geochim Cosmochim Acta 69:3967–3977.

81. Jones AM, Collins RN, Rose J, Waite TD. 2009. The effect of silica and natural organic matter on the Fe(II)-catalysed transformation and reactivity of Fe(III) minerals. Geochim Cosmochim Acta 73:4409–4422.

82. Lovley D. 2013. Dissimilatory Fe(III)- and Mn(IV)-Reducing Prokaryotes, p. 287–308. In Rosenberg, E, DeLong, EF, Lory, S, Stackebrandt, E, Thompson, F (eds.), The Prokaryotes. Springer Berlin Heidelberg, Berlin, Heidelberg.

83. Lovley DR, Phillips EJP. 1988. Novel mode of microbial energy metabolism: Organic carbon oxidation coupled to dissimilatory reduction of iron or manganese. Appl Environ Microbiol 54:1472–1480.

84. Straub KL, Benz M, Schink B. 2001. Iron metabolism in anoxic environments at near neutral pH. FEMS Microbiol Ecol 34:181–186.

85. Benz M, Schink B, Brune A. 1998. Humic acid reduction by *Propionibacterium freudenreichii* and other fermenting bacteria. Appl Environ Microbiol 64:4507–4512.

86. Kersting AB. 2013. Plutonium transport in the environment. Inorg Chem 52:3533–3546.

87. Kersting AB, Efurd DW, Finnegan DL, Rokop DJ, Smith DK, Thompson JL. 1999. Migration of plutonium in ground water at the Nevada Test Site. Nature 397:56–59.

88. Santschi PH, Roberts KA, Guo L. 2002. Organic nature of colloidal actinides transported in surface water environments. Environ Sci Technol 36:3711–3719.

89. Novikov AP, Kalmykov SN, Utsunomiya S, Ewing RC, Horreard F, Merkulov A, Clark SB, Tkachev VV, Myasoedov BF. 2006. Colloid transport of plutonium in the far-field of the Mayak Production Association, Russia. Science 314:638–641.

90. Schäfer T, Artinger R, Dardenne K, Bauer A, Schuessler W, Kim JI. 2003. Colloid-borne americium migration in Gorleben groundwater: Significance of iron secondary phase transformation. Environ Sci Technol 37:1528–1534.

91. Lujanien G, Beneš P, Štamberg K, Ščiglo T. 2012. Kinetics of plutonium and americium sorption to natural clay. J Environ Radioact 108:41–49.

